# Hydrogen production by *Sulfurospirillum* spp. enables syntrophic interactions of Epsilonproteobacteria

**DOI:** 10.1101/238212

**Authors:** Stefan Kruse, Tobias Goris, Martin Westermann, Lorenz Adrian, Gabriele Diekert

**Affiliations:** Department of Applied and Ecological Microbiology, Institute of Microbiology, Friedrich Schiller University, Philosophenweg 12, 07743 Jena, Germany; Center for Electron Microscopy of the University Hospital Jena, Ziegelmühlenweg 1, 07743 Jena, Germany; Department Isotope Biogeochemistry, Helmholtz Centre for Environmental Research – UFZ, Permoserstr. 15, 04318 Leipzig, Germany; Technische Universität Berlin, Fachgebiet Geobiotechnologie, Ackerstraße 76, 13355 Berlin

## Abstract

Hydrogen-producing bacteria are of environmental and biotechnological importance in anoxic environments, since hydrogen is an important electron donor for prokaryotes and of interest as an alternative energy source. Epsilonproteobacteria, inhabiting ecologically, clinically or biotechnologically relevant environments, are currently considered to be hydrogen-oxidizing bacteria exclusively. Here, we report hydrogen production for a genus of free-living Epsilonproteobacteria, *Sulfurospirillum* spp. inhabiting sediments, wastewater plants, bioelectrodes, oil reservoirs, contaminated areas, or marine habitats. The amount of hydrogen production was largely different in two subgroups of *Sulfurospirillum* spp., represented by *S. cavolei* and *S. multivorans*. The former is shown to be the more potent hydrogen producer and excretes acetate as sole organic acid, while the latter exhibited a more flexible fermentation, producing additionally lactate and succinate. The observed hydrogen production could be assigned to a group 4 hydrogenase similar to Hydrogenase 4 (Hyf) in *E. coli*. We propose that *Sulfurospirillum* spp. produce molecular hydrogen with electrons derived from pyruvate oxidation by pyruvate:ferredoxin oxidoreductase and reduced ferredoxin. This hypothesis is supported by comparative proteome data, in which both PFOR and ferredoxin as well as hydrogenase 4 are up-regulated. A co-culture experiment with *S. multivorans* and *Methanococcus voltae* cultivated with lactate as sole substrate shows a syntrophic interaction between both organisms, since the former cannot grow fermentatively on lactate alone and the latter relies on hydrogen as electron donor. This opens up new perspectives on microbial communities, since Epsilonproteobacteria could play a yet unrecognized role as hydrogen producers in anoxic microbial communities.

## Introduction

Hydrogen gas (H_2_), an important energy substrate for many bacteria and archaea, plays a crucial role in the anaerobic food web, e.g. in syntrophic interactions. It is produced by fermenting bacteria as a result of the disposal of excess reducing equivalents. Other prokaryotes may use it as an electron donor for e.g. sulfate respiration or methanogenesis. In syntrophic interactions, the H_2_-producing bacterium is dependent on the H_2_ uptake of its syntrophic partner, which sustains a low H_2_ partial pressure and thus enables H_2_ production, which would otherwise thermodynamically be unfavorable^1–3^. For example, butyrate, propionate or acetate-oxidizing anaerobic bacteria that form H_2_ as fermentation product are dependent on H_2_-oxidizing microorganisms such as methanogenic archaea^4–6^. It was shown that the interspecies H_2_ transfer becomes more efficient when syntrophs and methanogens are in close physical contact^7,8^. The syntrophic degradation of propionate by a co-culture of *Pelotomaculum thermopropionicum* and *Methanothermobacter thermoautotrophicus* as well as ethanol degradation by *Geobacter sulfurreducens* and *Geobacter metallireducens* resulted in aggregate formation and cell-to-cell contact of the involved organisms^9,10^. In addition to the importance of H_2_ in microbial food webs, H_2_ is considered to be an alternative energy source and biohydrogen production by microorganisms is discussed as one way to generate environmentally compatible fuels^11^.

Epsilonproteobacteria are hitherto considered to be H_2_-consuming organisms exclusively and H_2_-oxidizing enzymes of only a few Epsilonproteobacteria are characterized so far, e.g. the membrane-bound uptake hydrogenases of *Helicobacter pylori* and *Wolinella succinogenes*^12,13^. H_2_ production has never been shown to be performed by any Epsilonproteobacterium so far, although in recent years several Epsilonproteobacteria, especially marine, deep vent-inhabiting species, were reported to encode putative H_2_-evolving hydrogenases in their genomes^14–22^. *Sulfurospirillum* spp. are free-living, metabolically versatile Epsilonproteobacteria, many of which are known for their ability to respire toxic or environmentally harmful compounds such as arsenate, selenate or organohalides (e.g. tetrachloroethene - PCE)^23,24^. The anaerobic respiration with PCE, leading to the formation of cis-1,2-dichloroethene (cDCE), was studied in detail in *S. multivorans* (formerly known as *Dehalospirillum multivorans)*^26, 27^. Several *Sulfurospirillum* spp. were found in contaminated sediments, wastewater plants, marine environments or biocathodes^19, 23, 27, 28^. The role of *Sulfurospirillum* in such environments is unclear.

In previous studies, four gene clusters, each encoding a [NiFe] hydrogenase, were found in the genome of *S. multivorans*^20^ and most other *Sulfurospirillum* spp.^23^. Two of these appear to be H_2_-producing, the other two are potential H_2_-uptake enzymes as deduced from sequence similarity to known hydrogenases. Of these four hydrogenases, one of each type, H_2_-oxidizing and H_2_-producing, were previously detected in *S. multivorans^26,29^. The* periplasmically oriented H_2_-oxidizing enzyme is very similar to the characterized *W. succinogenes* and *H. pylori* membrane-bound hydrogenase (MBH). It comprises three subunits, the large subunit, harboring the NiFe active site, a small subunit for electron transfer with three FeS clusters, and a membrane-integral cytochrome *b*. The putative H_2_-producing, cytoplasmically oriented enzyme (Hyf) is a large, complex enzyme with eight subunits, four of them membrane-integral. Regarding amino acid sequence and subunit architecture, this hydrogenase is similar to hydrogenase 4 of *E. coli*, part of a putative second formate hydrogen lyase (FHL). However, in *S. multivorans*, Hyf is unlikely to form an FHL complex since the corresponding gene cluster does not encode any formate-specific proteins as is the case for the FHL complexes in *E. coli* (Supplementary Figure 1).

Here, we show that several *Sulfurospirillum* spp. produce H_2_ upon pyruvate fermentation. *S. cavolei* was observed to produce more H_2_ than other *Sulfurospirillum* spp., which is caused by a different fermentation metabolism. To unravel the metabolism and the hydrogenase equipment of both organisms, label-free comparative proteomics was carried out. A co-culture experiment of *S. multivorans* with the methanogenic archaeon *Methanococcus voltae* revealed an interspecies H_2_ transfer between both organisms suggesting a hitherto undiscovered contribution of *Sulfurospirillum* spp. and other Epsilonproteobacteria to the microbial anaerobic food web as a H_2_ producer.

## Experimental Procedures

### Cultivation of bacteria

S. *multivorans* (DSMZ 12446) was cultivated under anaerobic conditions at 28°C in a defined mineral medium^30^ without vitamin B_12_ (cyanocobalamin). Pyruvate (40 mM) was used as electron donor and fumarate (40 mM) as electron acceptor. For fermentation experiments, all cultivations were performed with pyruvate (40 mM) or lactate (40 mM) as sole energy source in the absence of an electron acceptor and without yeast extract. Bacteria were grown in serum bottles with a ratio of aqueous to gas phase of 1:1. If not stated otherwise, the gas phase was N_2_ (150 kPa). For the cultivation with 100% H_2_ in the gas phase, nitrogen was completely removed after autoclaving by flushing with H_2_ and an overpressure of 50 kPa was applied. Fermentation balance experiments were performed at 28°C in 1 L Schott bottles placed in a Fermentation apparatus to allow for the expansion of the gases during the cultivation and to determine the stoichiometry of dissolved and gaseous fermentation products (Supplementary Figure 2). For CO_2_ quantification, the gas phase of the Schott bottle was connected via a tube to a washing flask filled with 200 mL 4 M KOH to bind produced CO_2_ as carbonate. Downstream, the gas phase of the washing flask was further connected to a water-filled measuring cylinder placed up-side down in a water bath. The amount of H_2_ was determined volumetrically viathe displaced volume of water in the measuring cylinder that correlates with the amount of H_2_ produced. The concentration was calculated using the ideal gas equation. The adaptation experiment included a transfer in the next sub-cultivation step every 48 h with 10% inoculum. *Clostridium pasteurianum* W5 was cultivated in anoxic media composed of 1L basal medium (autoclaved) supplemented with the following anoxic solutions: 100 mL phosphate buffer (142 g L^−1^ K_2_HPO_4_, 15 g L^−1^ KH_2_PO_4_) and 5 mL iron solution (10 g L^−1^ FeSO_4_ · 7 H_2_O). The basal medium contained per L 142 mg NaCl, 1.42 g NH_4_Cl, 284 mg MgSO_4_· 7 H_2_O, 14.2 mg Na_2_MoO_4_ · 2 H_2_O, 28.4 mg D(+) biotin and 1.42 mg 4-aminobenzoate. Cells were grown in rubber-stoppered serum bottles with a ratio of aqueous to gas phase of 1:4. Pyruvate (40 mM) and Glucose (20 mM) were used as substrates. *Desulfitobacterium hafniense* DCB-2^32^ and *E. coli* JM109 were cultivated in medium described previously. The medium composition of the co-culture of *S. multivorans* and *Methanococcus voltae* DSMZ 1537 was identical to that described by Whitman *et al*.^32^, except that 5 g L^−1^ NaCl were added. Electron donor was 15 mM lactate. *C. pasteurianum* W5, *D. hafniense* DCB-2 and *E. coli* JM109 were taken from the strain collection of our laboratory and *M. voltae* was obtained from the German Collection of Microorganism (DSMZ, Braunschweig, Germany).

### Cell harvesting and preparation of cell suspensions and subcellular fractions

*S. multivorans, S. cavolei* and *C. pasteurianum* W5 cells were harvested in the mid-exponential growth phase in an anoxic glove box (COY, 134 Laboratory, Grass Lake, Michigan, USA) by centrifugation (12,000 x g, 10 min at 10°C). For the preparation of cell suspensions, the obtained cell pellets were washed twice in anoxic 100 mM MOPS-KOH-buffer (pH 7.0) and resuspended in two volumes (2 mL per g cells) of the same buffer. Subcellular fractionation was done by washing the cell pellet twice in 50 mM Tris-HCl (pH 8.0) and resuspension (2 mL per g cells) in the same buffer containing DNaseI (AppliChem, Darmstadt, Germany) and protease inhibitor (one tablet for 10 mL buffer; complete Mini, EDTA-free; Roche, Mannheim, Germany). The resuspended cells were disrupted using a beadmill (10 min at 25 Hz; MixerMill MM400, Retsch GmbH, Haan, Germany) with an equal volume of glass beads (0.25–0.5 mm diameter, Carl Roth GmbH, Karlsruhe, Germany). The crude extracts were separated from the glass beads by centrifugation (14,000 x g, 2 min) under anaerobic conditions and ultracentrifuged (36,000 x g, 45 min at 4°C). The obtained supernatants were considered as soluble fractions (SF). The pellets were washed twice with 50 mMTris-HCl (pH 8.0) including protease inhibitor and resuspended in the same buffer. The suspension was stated as membrane fraction (MF).

### Measurement of hydrogenase activity

H_2_ oxidizing activity was measured in H_2_-saturated buffer (50 mMTris-HCl, pH 8.0) with 1 mM benzyl viologen (BV) or methyl viologen (MV) at 30°C as artificial electron acceptors. The reduction of the redox dyes was followed at 578 nm using a Cary 100 spectrophotometer (Agilent Technologies, Waldbronn, Germany). H_2_-evolving activities of cell extracts were determined gas chromatographically with 1 mM methyl viologen as electron donor: MV was reduced with 20 mM sodium dithionite in an anoxic buffer system (50 mMTris-HCl, pH 8.0). Protein concentration was determined according to the method of Bradford^33^. Hydrogenase enzyme activities are given in nanokatal units (1 nmol H_2_ evolved per second).

### Analytical methods

Liquid samples were taken anaerobically, filtered with 0.2 μm-syringe filters (MiniSart RC4, Sartorius, Göttingen, Germany) and acidified with concentrated H_2_SO_4_ (2.5 μL mL^−1^ sample volume). Organic acids were separated at 50°C on an AMINEX HPX-87H column (7.8 x 300 mm, BioRad, Munich, Germany) with a cation H guard pre-column using 5 mM H_2_SO_4_ as mobile phase at a flow rate of -1 0. 7 mL min^−1^. The injection volume was 20 μL per sample. All acids (e.g. pyruvate, acetate, lactate, succinate and fumarate) were monitored by their absorption at 210 nm. Retention times were compared to known standards and concentrations were calculated using calibration curves. H_2_ was measured gas chromatographically with 99.999% argon as the carrier gas using a thermal conductivity detector (AutoSystem, Perkin Elmer, Berlin, Germany). Samples for gas analysis were taken from the gas phase with gas-tight syringes (Hamilton, Bonaduz, Switzerland). Concentrations were calculated using calibration curves. CO_2_ formed during the cultivation was determined gravimetrically. To 15 mL of the solution of the CO_2_ trap 7.5 mL NH_4_Cl (1 M) and 15 mL BaCl_2_ (1 M) were added and the pH was adjusted to 9 with concentrated HCl (37%). After stirring for 2 h at room temperature, the precipitated barium carbonate was filtered with filter circles and dryed over night at 80°C.

### Field emission-scanning electron microscopy (FE-SEM)

Field emissionscanning electron microscopy (FE-SEM) was performed with co-cultures of *S. multivorans* and *Methanococcus voltae*. After incubation of 3 mL culture in 2.5% glutaraldehyde for 15 min, the cells were pre-fixed for 2 h on poly-L-lysin coated cover slides (12 mm, Fisher Scientific, Schwerte, Germany). Washing of cover slides was done using 0.1 M sodium cacodylate (pH 7.2) (>98% purity, Sigma Aldrich, Steinheim, Germany) for three times. Subsequently, cells were post-fixed with 1% osmium tetroxide in the same cacodylate buffer and dehydrated with different ethanol concentrations. Critical point drying was done in a Leica EM CPD200 Automated Critical Point Dryer (Leica, Wetzlar, Germany) and the samples were coated with 6 nm platinum in a BAL-TEC MED 020 Sputter Coating System (BAL-TEC, Balzers, Liechtenstein). They were visualized at different magnifications using a Zeiss-LEO 1530 Gemini field emission scanning electron microscope (Carl Zeiss, Oberkochen, Germany).

### Sample preparation, mass spectrometry and proteome data analysis

Protein concentration of extracted proteins was determined using a Bradford reagent (Bio-Rad, Munich, Germany) with bovine serum albumin as standard. For protein identifications 20 μg of crude extracts were first cleaned from cations and cell debris by running shortly into an SDS gel. For this, the gel was run at 13 mA until the proteins entered the separating gel at a depth of about 3-5 mm. Then the protein band was cut out, reduced, alkylated and proteolytically digested with trypsin (Promega, Madison, WI, USA) and subsequently desalted with C18 ZipTips as described^34^.

Mass spectrometry was performed using an Orbitrap Fusion (Thermo Fisher Scientific, Waltham, MA, USA) coupled to a TriVersa NanoMate (Advion, Ltd., Harlow, UK). 5 μL of the peptide solution were separated using a Dionex Ultimate 3000 nano-LC system (Dionex/Thermo Fisher Scientific, Idstein, Germany). A sample volume of 1 μL was loaded onto a trapping column (300 μm inner diameter, packed with 5 μm C18 particles, Thermo Scientific) and separated on 15 cm analytical column (Acclaim PepMap RSLC, 2 μm C18 particles, Thermo Scientific) at 35°C. Liquid chromatography was done with a constant flow of 300 nL min^−1^ with a mixture of solvent A (0.1% formic acid) and B (80% acetonitrile, 0.08% formic acid) in a linear 90 min gradient of 4% to 55% solvent B.

MS1 scans were measured with a cycle time of 3 s in the Orbitrap mass analyzer between 350 and 2,000 *m/z* at a resolution of 120,000, automatic gain control (AGC) target 4×10^5^, maximum injection time 50 ms. Data-dependent acquisition (DDA) was employed selecting for highly intense ions (>5×10^4^) and charge state between +2 and +7 with a precursor ion isolation windows of 1.6 *m/z*. Fragmentation was done via higher energy dissociation (HCD) at 30% energy, and also measured in the Orbitrap analyzer at a resolution of 120,000 with an AGC target of 5×10^4^ and a maximum injection time of 120 ms. Fragmentation events were done within the 3 s of cycle time until the next MS1 scan was done excluding the same mass (±10 ppm) for further precursor selection for 45 s.

Mass spectrometric data were analyzed with Proteome Discoverer 1.4 (Thermo Scientific) against the NCBI *S. multivorans* database (CP007201.1) with the search engines SequestHT and MS Amanda. Oxidation of methionine was set as dynamic, carbamidomethylation of cysteine as static modification; two missed cleavages were accepted, mass tolerance of MS1 and MS2 measurements were set to 5 ppm and 0.05 Da, respectively. A percolator false discovery rate (FDR) threshold of <0.01 was set for peptide identification. Label-free quantification of proteins was done with the area of the three most abundant peptides of each protein. The values were logarithmized (log10) and normalized (see Supplementary Dataset 1) and a two-tailed T-test was applied. Significance values (p-values) of <0.05 were considered to indicate statistical significance. Only proteins identified in at least 50% of the three replicates (n≥2) were used for quantification, otherwise, proteins were considered to be identified.

## Results

### 1. Adaptation of *Sulfurospirillum multivorans* to pyruvate-fermenting conditions

In previous studies, *S. multivorans* and other *Sulfurospirillum* spp. were shown to grow fermentatively on pyruvate^23, 30, 35^. Only few data on growth behavior are available in the literature, but *S. multivorans* was reported to exhibit poor growth on pyruvate as sole energy source compared to respiratory growth with fumarate or tetrachloroethene (PCE) as electron acceptor^30^. However, we observed an adaptation of *S. multivorans* to fermentative growth on pyruvate. After about twenty transfers, a growth rate of 0.09 h^−1^ was determined (growth rate on pyruvate/fumarate, 0.19 h^−1^, Figure 1). During the adaptation to pyruvate fermentation, the growth rate increased on average by 0.02 h^−1^ with each transfer (Supplementary Figure 3). In addition, the lag phase duration decreased from initially 40h to 5h. After 18 transfers, no further significant increase of the growth rate was observed. This adaptation process was also observed for *S. cavolei, S. delyianum* and *S. arsenophilum*. For *S. barnesii* and *S. halorespirans*, no growth on pyruvate alone was detected, even after several subcultivation steps.

**Figure 1:**
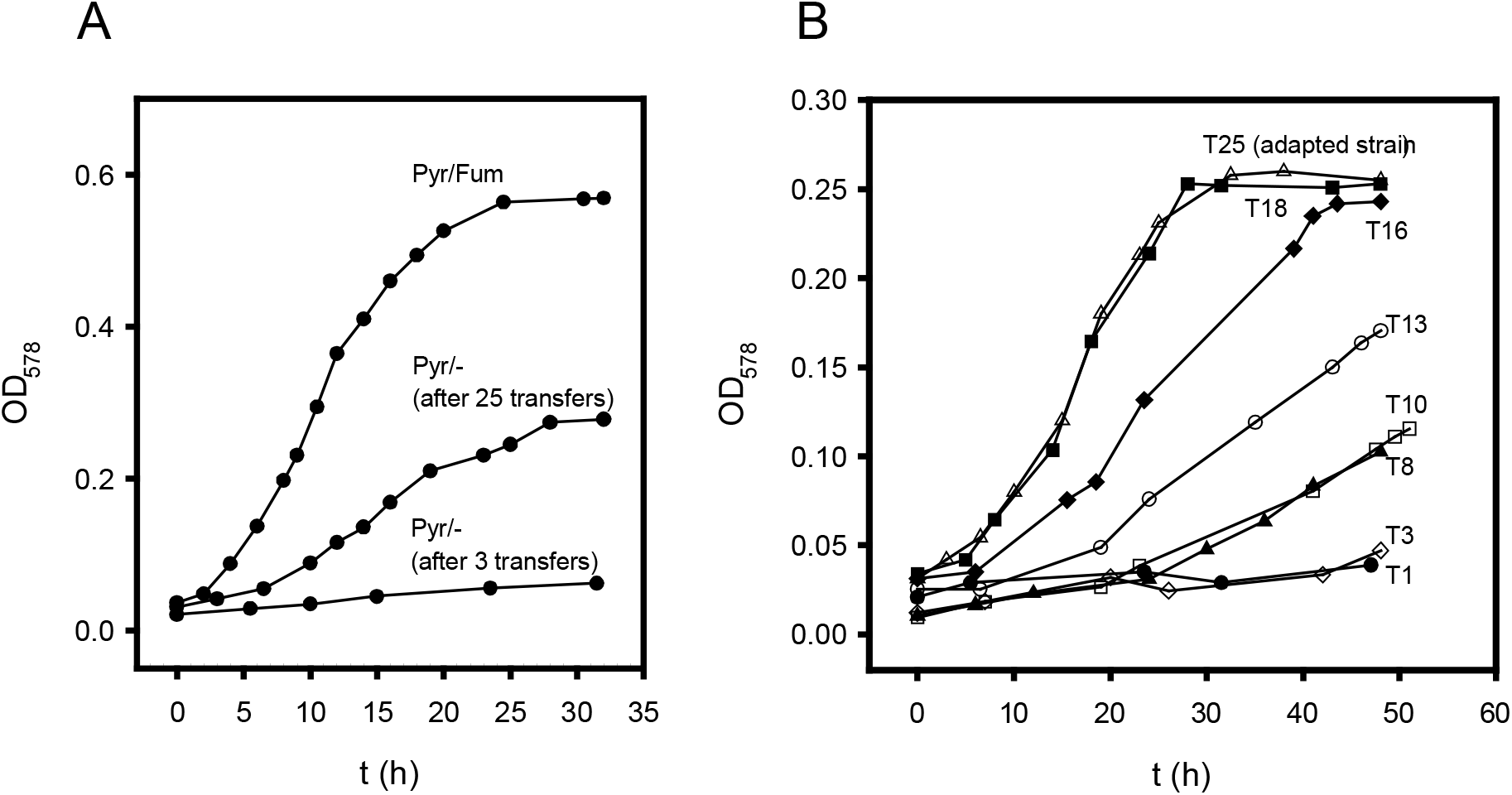
Adaptation of *S. multivorans* to pyruvate-fermenting conditions. A) Growth curves with pyruvate as sole growth substrate after three and twenty-five transfers; A culture with pyruvate/fumarate after three transfers is shown for comparison. B) Growth during continuous transfer on pyruvate without electron acceptor. Each transfer (10% inoculum) was done after 48 hours cultivation. Data were obtained from at least two independent biological replicates and are representatives. T - number of transfer step, Pyr - pyruvate, Fum - fumarate, OD_578_ - optical density at 578 nm.

### 2. Fermentative growth and H_2_ production of *Sulfurospirillum* spp

To get deeper insight into the fermentation pathways and H_2_ production capabilities of *Sulfurospirillum* spp., several species were cultivated with pyruvate as sole substrate. Six species were tested for pyruvate fermentation, of which *S. barnesii* and *S. halorespirans* were not able to grow even after cultivation for several months (data not shown). *S. cavolei, S. deleyianum* and *S. arsenophilum* were able to grow on pyruvate alone, albeit at slower rates than *S. multivorans* (0.03 h^−1^, 0.06 h^−1^, or 0.004 h^−1^, respectively). H_2_ production was measured for all fermentatively growing *Sulfurospirillum* spp., but the produced amount differed, depending on the species. *S. cavolei* produced the highest amount of H_2_ followed by *S. arsenophilum*. *S. deleyianum* and *S. multivorans* produced about 100 μmol per 100 mL culture. *D. hafniense* DCB-2, a known pyruvate-fermenting organohalide-respiring bacterium grows similar to *Sulfurospirillum* spp. (Figure 2A) but produced only minor amounts of H_2_ (20 μmol) (Figure 2b). Fermentative growth on lactate was not observed for any of the organisms including *D. hafniense* DCB-2 even after cultivation for several months (data not shown).

**Figure 2:**
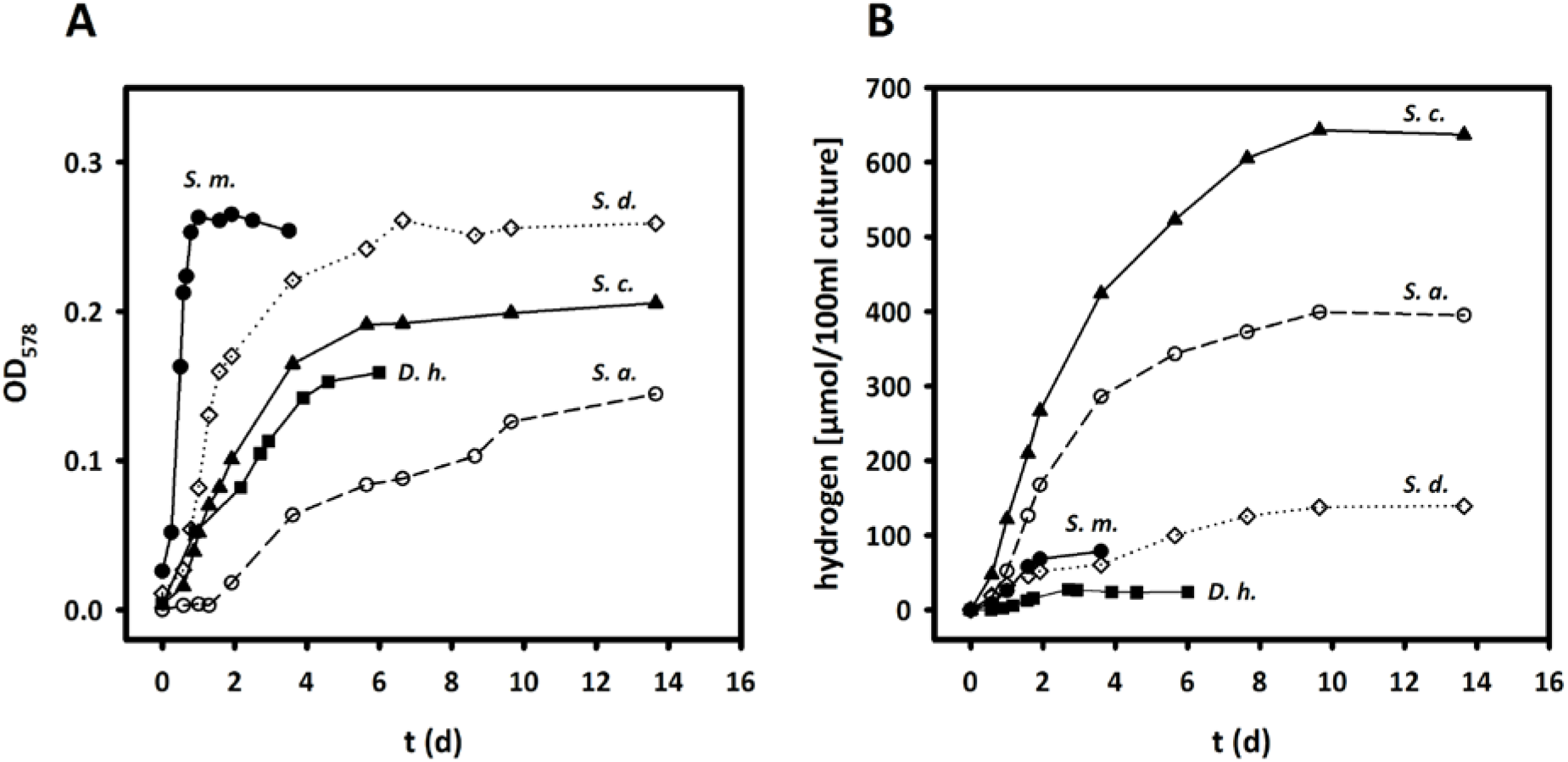
Growth (A) and H_2_ production (B) of *Sulfurospirillum* spp. and *D. hafniense* strain DCB-2 during fermentative growth on pyruvate after adaptation. The graph is a representative of three independent replicates. *S.m. - S. multivorans, S.d. - S. deleyianum, S.c. - S. cavolei, S.a. - S. arsenophilum, D.h. - D. hafniense* DCB-2.

### 3. Fermentative metabolism of *S. multivorans* and *S. cavolei*

To unravel the fermentative metabolism of two *Sulfurospirillum* spp. showing a different H_2_ production pattern during growth on pyruvate, *S. multivorans* and *S. cavolei* were cultivated in a fermentation apparatus in which the gas phase of the Schott bottle was connected to CO_2_ and H_2_ traps (see Supplementary Figure 2) to avoid increasing gas partial pressures and hence a possible product inhibition on H_2_ production or growth (see also next chapter). Fermentation products and pyruvate consumption were monitored via HPLC, GC, volumetric and gravimetric measurements in order to calculate the fermentation balance. In this experimental set up, a largely enhanced H_2_ evolution was measured when compared to the serum bottle experiment, with up to hundred times more H_2_ produced, while the growth was slower than in the previous set up (Figure 3A). After consumption of 40 mM pyruvate, 27 mM acetate, 10 mM lactate, 3 mM succinate,10 mM H_2_ and 28 mM CO_2_ were measured as fermentation products of *S. multivorans* (Figure 3A). *S. cavolei* showed slower growth than *S. multivorans* and a much higher amount of H_2_ evolved. During growth, which took 8 to 10 days, pyruvate (40 mM) was used up completely and 38 mM acetate, 36 mM H_2_ and 38 mM CO_2_ were the only products detected (Figure 3B). *S. deleyianum* showed similar fermentation products to *S. multivorans* (Supplementary Figure 4). The stoichiometry of the fermentation was verified by calculating the carbon recovery and an oxidation/reduction balance (Supplementary Table 1, Eqns (I) and (II)). In *S. multivorans*, the amount of reducing equivalents generated from pyruvate oxidation was calculated to be 54 [H], which fits to the amount of used reducing equivalents for the production of molecular hydrogen, lactate and succinate (52 [H], Supplementary Table 1). In *S. cavolei*, pyruvate oxidation leads to the generation of 76 [H], which were almost exclusively (72 [H]) used for proton reduction to H_2_. In addition, the carbon recovery is in agreement with the theoretical values and is 102.5% for *S. multivorans* and 95% for *S. cavolei*. The anabolic assimilation of the carbon source could be neglected due to the low amount of biomass produced.

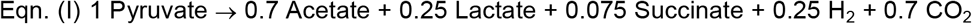

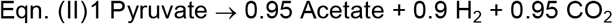

**Figure 3:**
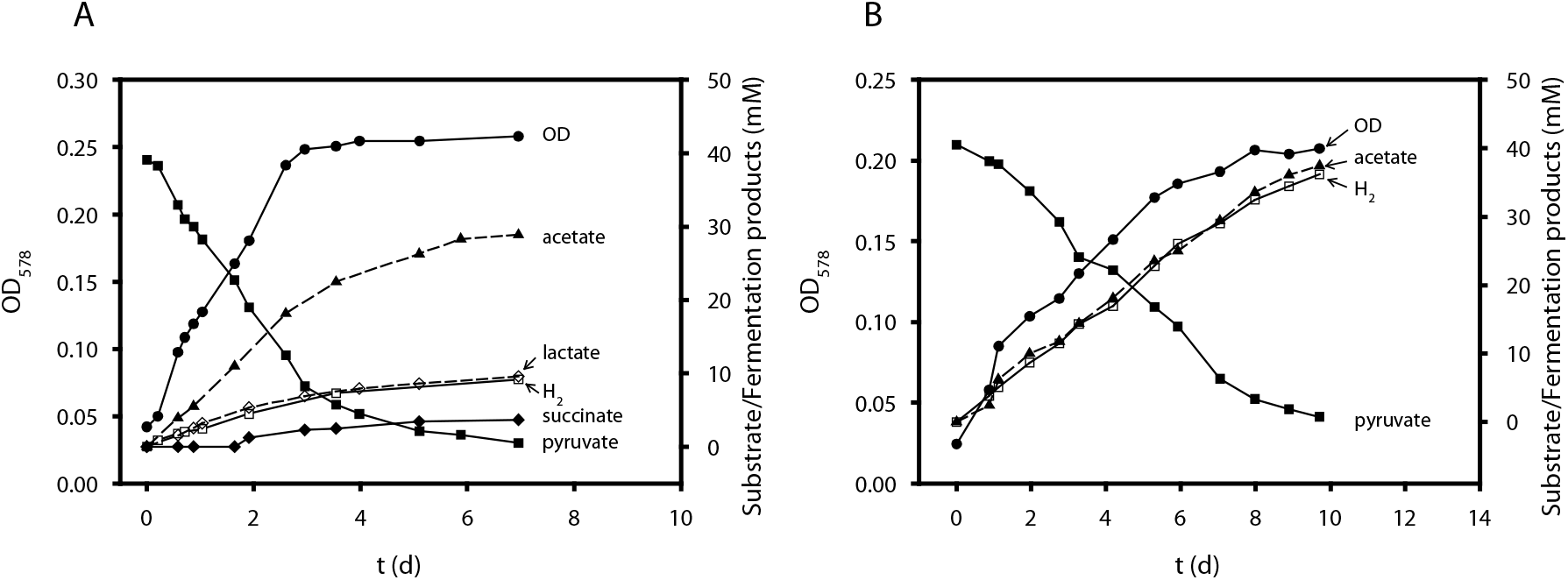
Fermentation balance of *S. multivorans* (A) and *S. cavolei* (B) during fermentative growth on pyruvate. Organic acids were measured via HLPC and h_2_ was determined volumetrically (for details see Materials and Methods).

### 4. Product inhibition by H_2_ on fermentative growth in *S. cavolei* and *S. multivorans*

The different amount of H_2_ produced in the growth experiments in serum bottles and the fermentation apparatus imply a product inhibition of H_2_ on H_2_ production. To investigate the effect of H_2_ in the gas phase on the fermentative growth of *S. multivorans* and *S. cavolei*, both organisms were cultivated in serum bottles with a gas phase of 100% H_2_ or 100% nitrogen (Figure 4). With nitrogen as gas phase, *S. multivorans* and *S. cavolei* showed similar growth and production rates of organic acids as observed in the fermentation apparatus. A strong negative effect on growth was observed with 100% H_2_ in the gas phase. *S. multivorans* was still able to ferment pyruvate but showed an inhibited growth and a lower cell density compared to the culture without H_2_ in gas phase, while *S. cavolei* was almost completely inhibited (Figure 4A). The restricted growth is also reflected by a lower pyruvate consumption rate (Figure 4B). In addition, the formation of fermentation products shifted from acetate production to lactate and succinate formation in *S. multivorans* (Figure 4C-E). *S. cavolei* produced neither lactate nor succinate and only minor amounts of acetate.

**Figure 4:**
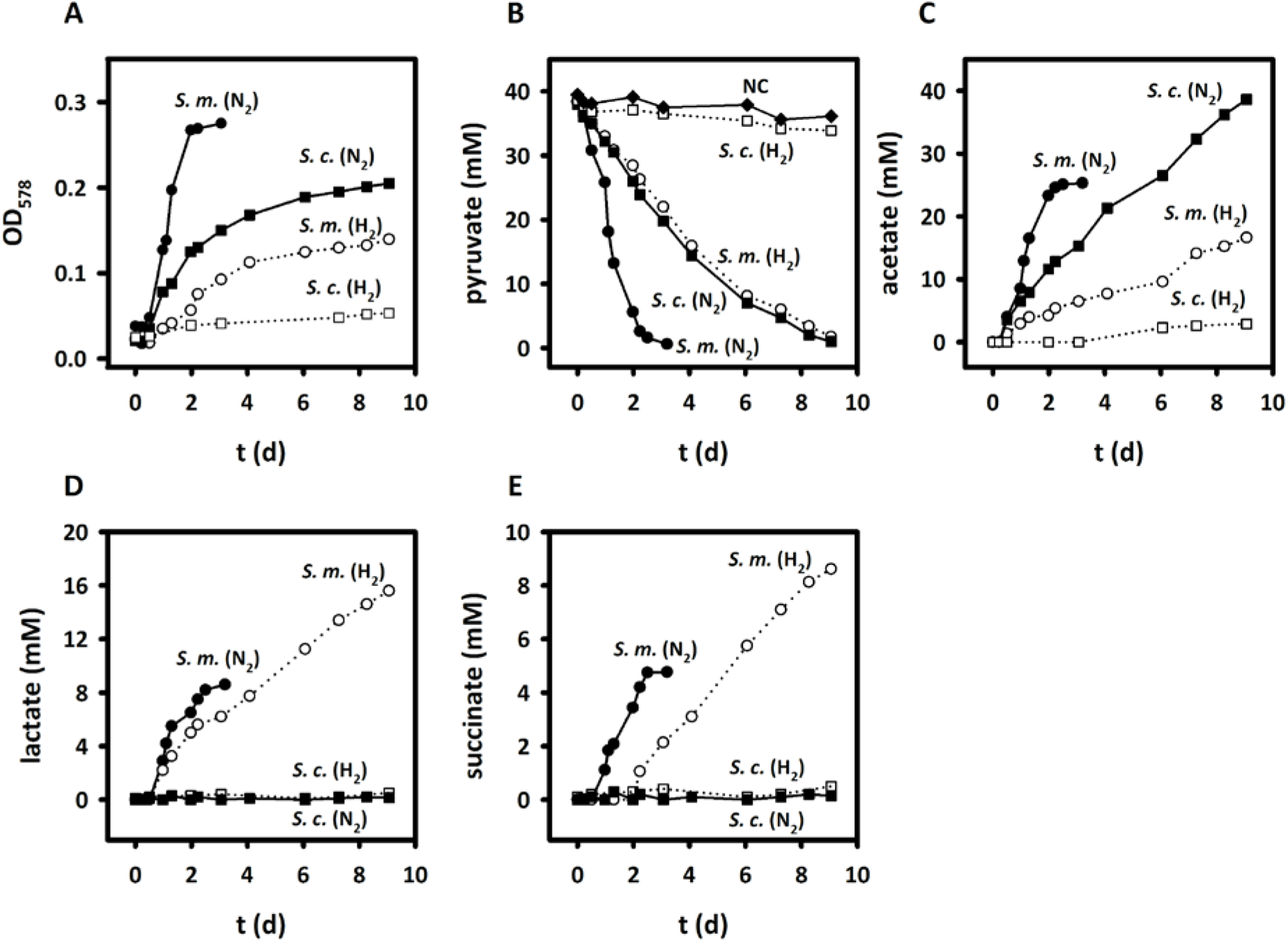
Growth and formation of fermentation products during cultivation under 100% nitrogen (N_2_) and 100% H_2_ atmosphere with pyruvate as sole energy source. Growth curve (A), pyruvate consumption (B) and acetate (C), lactate (D) and succinate (E) production are shown. Organic acids were measured via HPLC. Each cultivation was conducted in three biological replicates. S.m. - *S. multivorans*, S.c. - *S. cavolei*, N_2_ - nitrogen, H_2_ - hydrogen, NC - negative control (cell-free medium).

### 5. Hydrogenase activities by cell suspensions of *Sulfurospirillum* spp

The H_2_ production and oxidation capability of cell suspensions of *S. multivorans* and *S. cavolei* was analyzed to obtain further evidence about the hydrogenase involved in the production and oxidation reaction. Transcriptional and proteomic studies revealed the presence of two [NiFe] hydrogenases in *S. multivorans*^29^: a hydrogen-oxidizing periplasmic membrane-bound hydrogenase (MBH) and a putative H_2_-producing cytoplasmic membrane-bound hydrogenase (Hyf). These two hydrogenases might be distinguished by their different subcellular localization in hydrogenase acitivity assays. Photometrically measured H_2_-oxidizing activity was detected in whole cell suspensions as well as in membrane and soluble fractions (Table 1). In contrast, H_2_-producing activity, as monitored by GC, was only measured with membrane fractions of *S. multivorans* and *S. cavolei* with approximately 1.5-fold higher activity in the latter one. This suggests a cytoplasmic orientation of a membrane-associated H_2_-evolving hydrogenase, since methyl viologen should not have access to the cytoplasm of whole cells (Table 1). The H_2_-oxidizing activity of intact cells points towards a catalytic subunit accessible to benzyl and methyl viologen and, thus, a periplasmic orientation of the H_2_ uptake system. The membrane fractions of *S. multivorans* and *S. cavolei* cells grown on pyruvate as sole energy source were about 2-fold more active in H_2_-production than those of cells cultivated under respiratory growth conditions with pyruvate plus fumarate, while the latter exhibited slightly more H_2_ oxidation activity. *Clostridium pasteurianum* W5, which is known to harbor a soluble H_2_-producing hydrogenase, exhibited hydrogenase activity only in the soluble fractions and showed no H_2_ producing activity in cell suspensions with metyhl viologen as electron donor (Supplementary Table 2), thus serving as a control for the hydrogenase localization experiment.

**Table 1:** Hydrogen-production and oxidizing activities of cell suspensions and subcellular fractions of *S. multivorans* and *S. cavolei* cultivated under different growth conditions. Data are derived from three independent biological replicates.

| Cellular fraction |  | Hydrogenase activity (nkat mg<sup>-1</sup>) |  |  |  |  |  |
| --- | --- | --- | --- | --- | --- | --- | --- |
|  |  | *S. multivorans* |  |  | *S. cavolei* |  |  |
|  |  | MV → H<sub>2</sub> | H<sub>2</sub> → BV | H<sub>2</sub> → MV | MV → H<sub>2</sub> | H<sub>2</sub> → BV | H<sub>2</sub> → MV |
| **Cell suspensions** |  |  |  |  |  |  |  |
|  | Pyr | < 0.01 | 4.1 ± 0.5 | 0.7 ± 0.3 | < 0.01 | 5.5 ± 0.6 | 1.4 ± 0.3 |
| **Membrane fractions** |  |  |  |  |  |  |  |
|  | Pyr | 12.3 ± 2.4 | 23.5 ± 2.1 | n.d. | 20.6 ± 3.7 | 10.1 ± 0.5 | n.d. |
|  | Pyr + Fum | 5.7 ± 1.5 | 36.6 ± 3.3 | n.d. | 10.1 ± 2.4 | 13.5 ± 1.6 | n.d. |
| **Soluble fractions** |  |  |  |  |  |  |  |
|  | Pyr | < 0.01 | n.d. | n.d. | < 0.01 | n.d. | n.d. |
|  | Pyr + Fum | < 0.01 | 2.3 ± 0.3 | n.d. | < 0.01 | 1.9 ± 0.3 | n.d. |

MV → H<sub>2</sub> indicates H<sub>2</sub> formation activity, H<sub>2</sub> → BV/MV indicates H<sub>2</sub> oxidation. MV - methyl viologen, BV - benzyl viologen. Pyr - pyruvate, Fum - fumarate, n.d. - not determined

### 6. Comparative genomics and proteomics

To unravel the cause of the different fermentative metabolism of the two *Sulfurospirillum* sp., a comparative genomic analysis was done with the RAST sequence comparison tool^36^. Additionally, proteomes of *S. cavolei* NRBC109482 and *S. multivorans* cultivated under fermenting and respiring conditions with fumarate as electron acceptor were analyzed. Bidirectional blast hits with more than 50% amino acid sequence identity were considered as orthologs, proteins putatively fulfilling the same functions in both organisms. The genomes were overall similar, with 2057 of 2768 of the encoded proteins in *S. cavolei* being orthologs. Only few of the non-orthologous proteins in *S. cavolei* could be considered to play a role in the fermentation. Among the proteins encoded in the *S. cavolei* genome (annotated RefSeq WGS accession number NZ_AP014724), which do not have an ortholog in *S. multivorans*, we found a cluster encoding an [FeFe] hydrogenase known to contribute to fermentative H_2_ production in many bacteria, e.g. Clostridia (Supplementary Figure 5). A nearly identical gene cluster is found in the other two genomes of *S. cavolei* strains UCH003 and MES, the latter of which was assembled from a metagenome^19^. The large hydrogenase subunit gene, *hydA*, is disrupted by a stop codon resulting from a nucleotide insertion only in *S. cavolei* strain NRBC109482. The mutation was confirmed by PCR and Sanger sequencing. Transcript analysis of *hydA* suggested that mRNA of the [FeFe] hydrogenase active subunit gene was synthesized under pyruvate-fermenting growth conditions (Supplementary Figure 6). However, the [FeFe] hydrogenase was not identified in the proteome of *S. cavolei*.

Of the proteins related to pyruvate metabolism, a pyruvate, water dikinase (phosphoenolpyruvate [PEP] synthetase) is encoded in the genome of *S. multivorans* (encoded by SMUL_1602), but not in *S. cavolei*. This enzyme is responsible for the ATP-dependent synthesis of phosphoenolpyruvate from pyruvate in gluconeogenesis. The PEP synthetase was found in 6.3-fold higher abundance (p-value 0. 02) in the proteome of fermentatively cultivated *S. multivorans* cells (Supplementary Table 3). In *S. cavolei*, PEP might be formed from pyruvate via oxaloacetate by two reactions catalyzed by pyruvate carboxylase and PEP carboxykinase. These two enzymes are encoded in one gene cluster (SCA02S_RS02520 and SCA02S_RS02525, respectively, Supplementary Figure 7). In *S. multivorans* these proteins (SMUL_0789 and SMUL_0791) cluster with a gene encoding a subunit similar to the membrane subunit of a putative Na^+^-translocating oxaloacetate decarboxylase (SMUL_0790), of which an ortholog is not encoded in *S. cavolei* (Supplementary Figure 7). Both pyruvate carboxylase/oxaloacetate decarboxylase and PEP carboxykinase were found in the proteomes of both organisms in slightly higher amounts in cells grown with pyruvate only (Supplementary Dataset 2). Interestingly, also *S. arsenophilum*, producing larger amounts of H_2_ than *S. multivorans* (Figure 2), lacks the putative oxaloacetate decarboxylase subunit gene (Supplementary Figure 7).

The Hyf hydrogenase was found in high abundancies especially in the proteome of *S. multivorans* cultivated with pyruvate alone. Here, four out of eight of the structural subunits were found in the 10% of the most abundant proteins, while none were found in the top 10% under respiratory conditions. In *S. cavolei*, the hydrogenase-4 subunits were not as abundant as in *S. multivorans* with only two out of six quantified subunits in the top 20% (Supplementary Dataset 2). In both organisms, a significantly higher amount of Hyf subunits was quantified under fermentative growth conditions (S. *multivorans:* 4- to 27-fold for the structural subunits HyfA-HyfI, all p-values are <0.001, *S. cavolei:* 2- to 5-fold for HyfA-HyfI, all p-values are <0.05; Figure 5, Supplementary Table 3, Supplementary Dataset 2). Interestingly, the Hyf gene cluster is disrupted at one site in *S. halorespirans*, which cannot grow on pyruvate alone. Genome sequencing^37^ revealed a transposase insertion at *hyfB* which might result in a non-functional gene *S. halorespirans*. The transposon insertion was confirmed by PCR using *hyfB-* specific primers flanking the transposase (Supplementary Figure 9). The membrane-bound subunits HyfE and HyfF were found in fermenting cells of *S. multivorans* exclusively. Sequence comparison of the Hyf-like hydrogenase of *S. multivorans* shows similarities to the formate hydrogen lyase complex of *E. coli* and to complex I of *Thermus thermophilus* (Supplementary Figure 8). An analysis of the potential proton-translocating sites in Hyf of *S. multivorans* and a comparison to the FHL of *E. coli* and the related subunits of several complex I is given in the Supplementary information (Supplementary Note 1, Supplementary Table 5 and Supplementary Figures 10 - 12).

**Figure 5:**
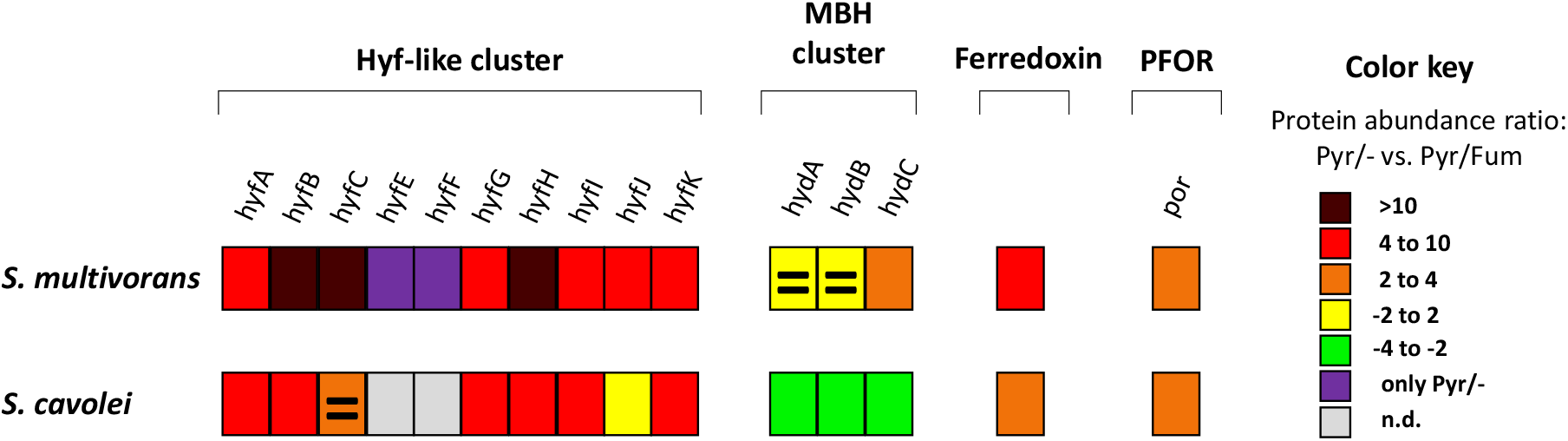
Comparative proteomics of proteins possibly involved in pyruvate fermentation of *S. multivorans* and *S. cavolei*. Comparison of cells grown with pyruvate alone (fermentative conditions) was done with cells grown with pyruvate/fumarate (standard = respiratory conditions). For quantified proteins the protein intensity is given as coloured squares. Non-significantly altered proteins are marked with an equal sign. Proteins exclusively found in pyruvate fermenting cells are colored light blue. All data were obtained from 3 independent biological replicates. Ratios in dashed boxes not significantly altered (p-values >0.05). Hyf-like - Hyf hydrogenase (SMUL_2383-2392; SCA02S_RS01920-RS01965), MBH - membrane-bound hydrogenase (SMUL_1423-1425; SCA02S_RS01350-RS01360), Fd - ferredoxin (SMUL_0303; SCA025_RS12260), PFOR - pyruvate:ferredoxin-oxidoreductase (SMUL_2630; SCA02S_RS04525). Pyr - pyruvate, Fum - fumarate.

**Figure 6:**
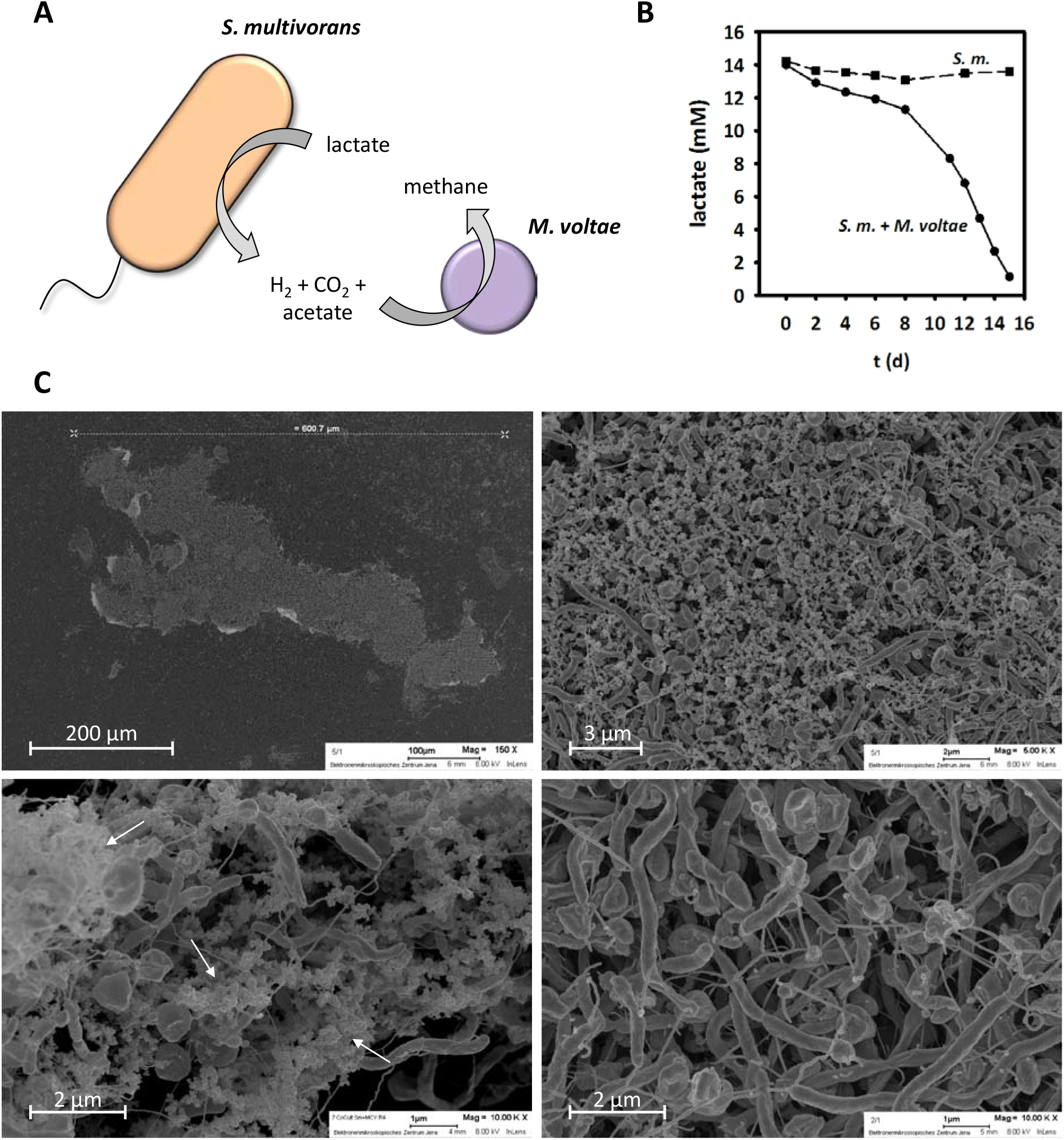
Syntrophic co-culture of *S. multivorans* and *Methanococcus voltae*. (A) scheme of syntrophic interactions and exchange of metabolites and (B) lactate concentration in *S. multivorans* pure culture and co-culture of *S. multivorans* and *M. voltae*. (C) Electron microscopic analyses and images of formed aggregates. Image sections were obtained from different areas of an aggregate. White arrows indicate EPS-like structures. Cultivation experiments included three biological replicates. S.m. - *S. multivorans*.

A search for the *hyf* gene cluster in genomes of Epsilonproteobacteria shows that it is ubiquitous in, but not limited to, *Sulfurospirillum* spp (Supplementary Table 4). Four out of 15 *Sulfurospirillum* sp. genomes harbor a second *hyf* gene cluster co-located with genes encoding a formate transporter and a formate dehydrogenase (Supplementary Figure 1). In *Arcobacter* spp. and the marine species *Caminibacter mediatlanticus* and *Lebetimonas* spp., only the latter gene cluster encoding a putative FHL complex is found. In several *Campylobacter* spp. including *C. concisus*, a *hyf* gene cluster with a formate transporter gene was identified (Supplementary Figure 1), while a second group of *Campylobacter* (including *C. fetus*) does not encode any formate-related proteins (Supplementary Table 4).

A pyruvate:ferredoxin oxidoreductase (PFOR) and a ferredoxin (Fd) showed also a higher abundance in both *Sulfurospirillum* sp. under fermenting conditions (S. *multivorans:* PFOR 2-fold, Fd 6-fold, *S. cavolei:* PFOR 4-fold, Fd 2-fold, all p-values are <0.01; Figure 5, Supplementary Table 3). A second pyruvate-oxidizing enzyme, a quinone-dependent pyruvate dehydrogenase encoded exclusively in the genome of *S. multivorans*, was significantly lower abundant during pyruvate fermentation (7-fold, p-value 0.02). The enzymes responsible for ATP generation via substrate-level phosphorylation, phosphotransacetylase and acetate kinase, are slightly higher abundant during pyruvate fermentation in both *Sulfurospirillum* sp. (approximately 2-fold for both enzymes in *S. multivorans*, p-values are <0.01 and approximately 3-fold in *S. cavolei*, p-values are <0.001; Figure 7, Supplementary Table 3). The malic enzyme is higher abundant during fermentation in *S. multivorans* (3.7-fold, p-value <0.001, Supplementary Table 3) and not quantified in any proteome of *S. cavolei* (Supplementary Table 3). The fumarate hydratase is found in lower abundancies during fermentation in both organisms (S. *multivorans:* 2-fold, p-values <0.01, *S. cavolei:* 6-fold, p-values <0.001), while the fumarate reductase is significantly lower abundant only in *S. cavolei (S. cavolei:* approximately 6-fold, p-values <0.001). The subunits of the membrane-bound hydrogenase (MBH) were quantified in either unsignificantly lower amounts (HydAB, approximately 2-fold, p-values 0.40 and 0.07) or slightly higher amounts (HydC, approximately 2-fold, p-value 0.01) under fermenting conditions for *S. multivorans*. In contrast, HydABC were found to in significantly lower amounts in *S. cavolei* when grown fermentatively. Of the cytoplasmic H_2_-producing hydrogenase (Ech-like), only one subunit was quantified in *S. multivorans* grown with pyruvate alone; this subunit was classified in the lower 50% abundant proteins. In *S. cavolei*, five of six Ech-like hydrogenase subunits were quantified in cells cultivated with pyruvate alone and two of six subunits in pyruvate/fumarate-cultivated cells, all of the Ech-like subunits were found in the lowest third abundant proteins. No subunit of the cytoplasmic uptake hydrogenase (HupSL) was found in any of the proteomes.

**Figure 7:**
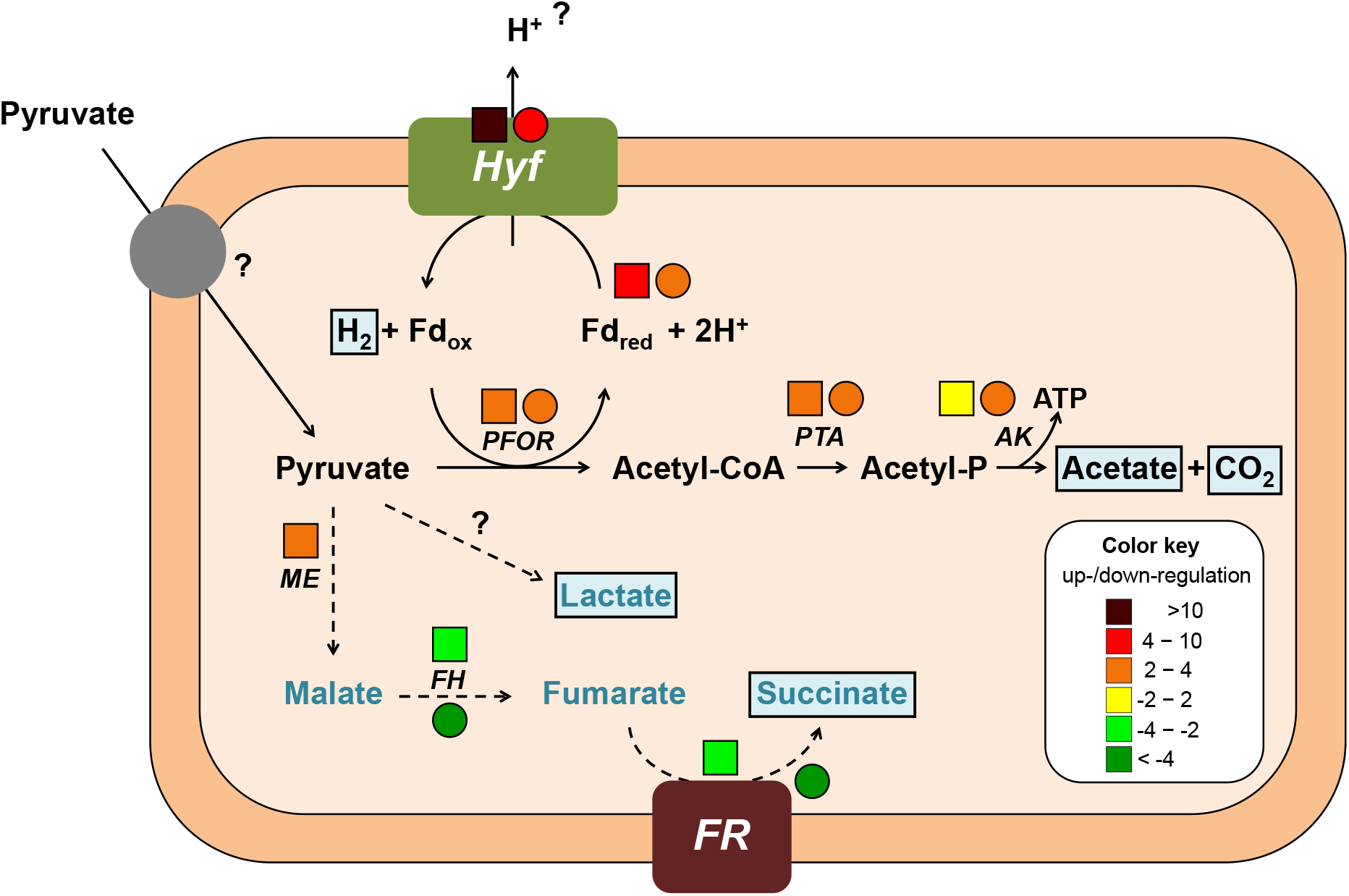
Tentative scheme of pyruvate fermentation metabolism in *S. multivorans* and *S. cavolei*. Reactions represented by solid arrows belong to the core pyruvate metabolism and are catalyzed by both organisms. Reactions with dashed arrows are solely catalyzed by *S. multivorans*, fumarate hydratase and fumarate reductase are also present in *S. cavolei*. Fermentation products are highlighted in light blue boxes. Protein abundance ratios (pyruvate alone versus pyruvate/fumarate) are indicated by colored squares (*S. multivorans*) and circles (*S. cavolei*) at the protein abbreviations (. Color code of the ratios is given in the box at the lower right. Hyf - Hyf-like hydrogenase (SMUL_2383-2392; SCA02S_RS01920-RS01965), PFOR - pyruvate:ferredoxin oxidoreductase (SMUL_2630; SCA02S_RS04525), PTA - phosphotransacetylase (SMUL_1483; SCA02S_RS00245), AK - acetate kinase (SMUL_1484; SCA02S_RS00240), ME - malic enzyme (SMUL_3158; corresponding enzyme in *S. cavolei* is not present), FH - fumarate hydratase (SMUL_1459, SMUL_1679-1680; SCA02S_RS00615-RS00620), FR - fumarate reductase (SMUL_0550-0552; SCA02S_RS07735-RS07740).

A putative lactate dehydrogenase (SMUL_0438, SCA02S_RS08360) with 35% amino acid sequence identity to a characterized lactate-producing lactate dehydrogenase from *Selenomonas ruminantium*^38^ was not detected in any proteome. This is in accordance to the lack of pyridine dinucleotide-dependent lactate-oxidizing or pyruvate-reducing activities in cell extracts of *S. multivorans* (data not shown, methods described in the Supplement). Several candidates for pyridine dinucleotide-independent lactate dehydrogenases (iLDH) are encoded in the genome of *S. multivorans*. Since *S. deleyianum* shows also lactate production during pyruvate fermentation, only genes present as orthologs in both genomes were considered to be responsible for lactate production in *Sulfurospirillum* spp. Functionally characterized iLDHs are flavin and FeS-cluster-containing oxidoreductases^40^ or enzymes related to malate:quinone oxidoreductase^40^. Only two candidates of the former class were identified in the genome, encoded by SMUL_1449 and SMUL_2229. Of these, only the latter gene product was detected in the proteome, however, not in altered amounts under fermentative conditions when compared to respiratory cultivation.

### 7. *Sulfurospirillum multivorans* as a syntrophic partner for *Methanococcous voltae*

To unravel the potential role of *S. multivorans* in a syntrophic partnership as H_2_ producer, a co-culture with *Methanococcus voltae* was prepared. *M. voltae* is a methanogenic archaeon dependent on H_2_ or formate as electron donor and CO_2_ as electron acceptor^32^. To investigate the syntrophic interaction of the two organisms, the co-culture was cultivated with lactate, which could not serve as a fermentation substrate for pure *S. multivorans* cultures. A syntrophic, hydrogen-consuming partner keeping H_2_ concentration at a low level in co-cultures might render lactate fermentation by *S. multivorans* thermodynamically feasible in a co-culture. In the corresponding co-culture, 15 mM lactate was consumed in approximately two weeks, indicating lactate fermentation by *S. multivorans* and H_2_ transfer to *M. voltae* as syntrophic partner (Figure 6 A,B). Formation of methane was measured gas-chromatographically (data not shown). Electron microscopic analyses of the co-culture revealed cell aggregates with sizes between 50 and 600 μm (Figure 6C, Supplementary Figure 13). These aggregates showed a compact network of the rod-shaped *S. multivorans* and coccoidal *M. voltae* with net-forming flagellum-like structures surrounding the organisms. The cells in the aggregates are embedded in extracellular polymeric substances (EPS)-like structures, which might aid cell-to-cell contact.

## Discussion

In this study, production of H_2_ was observed for several *Sulfurospirillum* species during pyruvate fermentation, which is the first evidence of H_2_ production for Epsilonproteobacteria, which hitherto were generally regarded as H_2_ oxidizers^14,35,41,42^. Specifically, we report H_2_ production for *S. multivorans, S. cavolei, S. arsenophilum* and *S. deleyianum* during fermentative growth on pyruvate. Sequential subcultivation on pyruvate alone revealed a continuous adaptation of *Sulfurospirillum* spp. to a fermentative metabolism. The mechanisms behind this long-term adaptation process in *Sulfurospirillum* spp. remain unresolved for now and might include genomic rearrangements and/or population dynamics, but also a long-term regulatory effect similar to the one observed for *S. multivorans* after continuous transfer without PCE as electron acceptor^43^ might play a role. The basis for the latter effect is also unknown to date.

Two different fermentation balances were observed for the different *Sulfurospirillum* spp. tested. While *S. cavolei* showed the highest H_2_ production rate and produced, besides hydrogen, acetate and CO_2_ exclusively, *S. deleyianum* and *S. multivorans*, displaying lower H_2_ production, additionally produced succinate and lactate. Pyruvate is most likely oxidized to acetate by the pyruvate:ferredoxin oxidoreductase, which showed an upregulation in the protome of fermentatively cultivated compared to fumarate-respiring cells in both, *S. multivorans* and *S. cavolei*. The quinone-dependent pyruvate dehydrogenase (PoxB) might transfer electrons generated upon pyruvate fermentation to menaquinone, however, since PoxB is downregulated in fermenting cells, it is suggested that this enzyme does not contribute significantly, if at all, to pyruvate oxidation under this condition. A pyruvate formate lyase is not encoded in any *Sulfurospirillum* spp., which, in addition to the low protein abundance of a cytoplasmic formate dehydrogenase, argues against the role of the Hyf in a formate hydrogen lyase complex as suggested for a similar hydrogenase in *E. coli*^44^. The generated acetyl-CoA is used to generate acetate and one mol ATP per mol pyruvate via substrate-level phosphorylation.

Electrons generated upon pyruvate oxidation are most likely transferred in both organisms to a ferredoxin of the *Allochromatium vinosum-type*, which is known for the very negative redox potentials of its two [4Fe4S] clusters^45^. The proteome data and biochemical experiments presented in our study strongly suggest that the Hyf (hydrogenase 4) of *Sulfurospirillum* spp. accepts electrons from the reduced ferredoxin to reduce two protons to hydrogen. Hyf is significantly upregulated, whereas the other hydrogenases are either detected only in low amounts in the proteome data or are unaltered or downregulated under fermentative cultivation. Furthermore, reduced methyl viologen served as electron donor for H_2_ production only with crude extract and not with intact cells, suggesting a cytoplasmic localization of the hydrogen-producing hydrogenase, as was suggested for the Hyf^20,29^. The involvement of a Hyf in H_2_ production via pyruvate oxidation was also observed in a group 4 hydrogenase from *Pyrococcus furiosus* ^46^ and for a genetically modified *E. coli* strain^47^. The structure and subunit composition of several group 4 hydrogenases suggested their involvement in the generation of a proton motive force, thereby contributing to ATP formation^48,49^. A thorough alignment analysis of the subunits of *Sulfurospirillum* spp. Hyf indicated that most of the important residues in the membrane helices are conserved, thus making a role in energy conservation of this hydrogenase a possible scenario. The difference in the amount of H_2_ produced with *S. cavolei* producing more H_2_ than *S. multivorans* can be explained by two different fermentation metabolism types. Opposed to *S. cavolei*, reducing equivalents can be channelled into the production of lactate and succinate by *S. multivorans* (as was also observed for *S. deleyianum)* upon pyruvate fermentation. Succinate might be produced from fumarate (fumarate reductase) via malate (fumarase), which could be formed from pyruvate via reductive decarboxylation to malate by the malic enzyme. This enzyme, which often functions in the reverse direction e. g. in C_4_ plants, is upregulated in *S. multivorans* under fermentative conditions. This finding supports the involvement of the malic enzyme in conversion of pyruvate to malate. The malic enzyme was not detected in the proteomes of *S. cavolei*, which might at least partially explain the different fermentation balances.

The origin of lactate in *S. multivorans* is not clear. An NAD^+^-dependent lactate dehydrogenase was not detected in any of the proteomes and no NAD(P)^+^-dependent lactate production could be measured. Most likely, the lactate dehydrogenase is misannotated in the genome of *S. multivorans*, as reported for a related protein of *Campylobacter jejuni*^50^. A possible source of lactate could be the reduction of pyruvate via an unknown, NAD^+^-independent lactate dehydrogenase (iLDH). Some of these are characterized to be functional in the direction of lactate oxidation^51,52^ and could act in the reverse direction to produce lactate in *Sulfurospirillum* spp., possibly with reduced ferredoxin as electron donor. Several candidates of iLDHs are encoded in the genome of *S. multivorans*, but only one of them shows a slight upregulation on pyruvate alone. A homolog of the corresponding gene cluster is not encoded in the lactate-producing *S. deleyianum*, making it an unlikely candidate for lactate production. A glycolate oxidase was shown to be responsible for lactate oxidation in *Pseudomonas putida*^39,53^ and a homolog is encoded in both lactate-producing *Sulfurospirillum* spp. This protein, however, is not upregulated upon pyruvate fermentation and further studies are needed to identify the lactate-producing enzyme in *S. multivorans*.

The different disposal of excess reducing equivalents during fermentation enables *S. multivorans* to grow with pyruvate even with 100% H_2_ in the gas phase, whereas the growth of *S. cavolei* was nearly completely abolished under these conditions. This correlates with a shift towards a higher production of lactate and succinate and a lower acetate and H_2_ production of *S. multivorans* under these conditions. The inability of *Sulfurospirillum* spp. to use lactate as sole substrate in pure cultures is most probably due to the thermodynamically unfavorable lactate oxidation to pyruvate upon hydrogen production. However, a syntrophic partnership of *S. multivorans* with a hydrogen-consumer, *Methanococcus voltae*, enabled lactate utilization by *S. multivorans* and led to the formation of large cell aggregates of the two organisms presumably via the formation of EPS.

These findings confirm our suggested role of *Sulfurospirillum* spp. as H_2_ producers in anaerobic food webs. Additionally, this role as a potential H_2_ producer is most likely not limited to this genus. In a genome mining approach, hyf gene clusters were found among several genera of Epsilonproteobacteria inhabiting a wide range of habitats. Some *Campylobacter* spp. known to be opportunistic or food-born pathogens encode the same Hyf as *Sulfurospirillum* spp., while hyf gene clusters containing either a formate channel gene in different *Campylobacter* spp. or additionally a cytoplasmic formate dehydrogenase in other phyla might indicate the formation of a formate hydrogen lyase complex. Since a PFL is missing in these bacteria, it might be presumed that extracellular formate might aid growth in these bacteria as reported for *Thermococcus* spp.^54^. Some *Sulfurospirillum* spp. even encode for both, a FHL-independent Hyf and one presumably forming an FHL complex, pointing towards seperate regulation and roles of both hydrogenases and thus for even more physiological diversity in this genus.

## Conclusion

Taken together, our results show that several Epsilonproteobacteria have to be considered as H_2_ producers and serve as syntrophic partners under certain conditions. H_2_ production in *Sulfurospirillum* spp. under the tested conditions relies on Hyf, a multisubunit, membrane-bound and cytoplasmically oriented group 4 NiFe hydrogenase similar to the one used in a second *E. coli* formate hydrogen lyase complex and probably functioning as a proton pump. Adaptation to fermentative conditions seems to be common in *S. multivorans* and related strains, although the underlying mechanism of this process is still unclear. Two seperate clades of *Sulfurospirillum* spp. have different fermentation pathways, the *S. cavolei* clade producing more H_2_ and exclusively one organic acid, namely acetate, in comparison to *S. multivorans*, which additionally produces lactate and succinate. All these findings imply an even higher versatility for Epsilonproteobacteria than previously thought and a new ecological role for *Sulfurospirillum* spp., which inhabit a large range of environmentally or biotechnologically important habitats such as wastewater plants, oil reservoirs, bioelectrodes, contaminated sediments or marine areas.

## Acknowledgement

This work was funded by the German Research Council (DFG) - Jena School for Microbial Communication (JSMC) and Research Unit FOR1530. We would like to gratefully acknowledge Susanne Linde (University Hospital Jena, Center for Electron Microscopy) for the field-emission scanning electron microscopic analysis. The presented work included the use of analytical facilities of the Centre for Chemical Microscopy (ProVIS) at the Helmholtz Centre for Environmental Research (UFZ Leipzig). ProVIS is funded by the European Regional Development Funds (EFRE - Europe funds Saxony) and the Helmholtz Association. The authors would like to thank Benjamin Scheer (UFZ Leipzig) for invaluable assistance in the lab with mass spectrometry and Dominique Türkowsky (UFZ Leipzig) for help with statistical analysis of proteome data.

## Author Contributions

SK performed the wetlab work, SK and TG planned experiments, TG initiated the study, TG and GD supervised the study, LA performed mass-spectrometric analysis, SK, TG and GD analyzed and discussed data, MW was responsible for electron microscopy, SK and TG drafted the manuscript, all authors revised, read and approved this manuscript.

